# Multicomponent regulation of actin barbed end assembly by twinfilin, formin and capping protein

**DOI:** 10.1101/2023.04.24.538010

**Authors:** Heidi Ulrichs, Ignas Gaska, Shashank Shekhar

## Abstract

Living cells assemble their actin networks by regulating reactions at the barbed end of actin filaments. Formins accelerate elongation, capping protein (CP) arrests growth and twinfilin promotes depolymerization at barbed ends. How cells integrate these disparate activities within a shared cytoplasm to produce diverse actin networks, each with distinct morphologies and finely tuned assembly kinetics, is unclear. We used microfluidics-assisted TIRF microscopy to investigate how formin mDia1, CP and twinfilin influence the elongation of actin filament barbed ends. We discovered that the three proteins can simultaneously bind a barbed end in a multiprotein complex. Three-color single molecule experiments showed that twinfilin cannot bind actin filament ends occupied by formin mDia1 unless CP is present. The trimeric complex is short-lived (∼1s) and results in rapid dissociation of CP by twinfilin causing resumption of rapid formin-based elongation. Thus, the depolymerase twinfilin acts as a pro-formin factor that promotes polymerization when both CP and formin are present. While a single twinfilin binding event is sufficient to displace CP from the trimeric complex, it takes about 30 independent twinfilin binding events to remove capping protein from CP-bound barbed end. Our findings establish a new paradigm in which polymerases, depolymerases and cappers work in concert to tune cellular actin assembly.

## Introduction

Cellular actin dynamics are essential in several key processes such as cell migration, wound healing, cell division and endocytosis^1, 2^. Depending upon requirements of specific processes, cells dynamically tune the rate of assembly, filament length, and internal architecture of filamentous actin networks. For example, while formins assemble fast-elongating linear actin networks comprising of long bundled filaments e. g. in filopodia, stereocilia and stress fibers^3–7^, the Arp 2/3 complex assembles dendritic actin arrays made of short, relatively slowly growing branched actin filaments e.g. in lamellipodia of motile cells and at sites of endocytosis^1, 8, 9^. How cells assemble actin networks with such diverse morphologies and dynamics in a shared cytoplasm remains an open question.

Intracellular actin assembly is primarily governed by reactions occurring at the barbed end of actin filaments (sometimes also referred to as the plus end)^2, 10^. Even though actin filaments can elongate by spontaneous addition of actin monomers, barbed end dynamics can be further tuned by three distinct classes of proteins which directly bind barbed ends. These proteins include polymerases, cappers and depolymerases. Polymerases, like formins and Ena/VASP, nucleate actin filaments and support filament elongation by remaining processively bound to filament barbed ends ^11, 12^. Formins can additionally accelerate the rate of barbed-end assembly by up to five-fold in presence of profilin-bound actin monomers (referred to as profilin-actin monomers or PA henceforth)^13, 14^. Cappers, like capping protein (CP) and gelsolin, cause complete arrest of filament growth by preventing monomer addition at free barbed ends ^15, 16^. Depolymerases, like twinfilin, comprise the third class of barbed-end binding proteins^17, 18^. Initially discovered as an actin monomer-sequestering protein^19^, twinfilin induces barbed-end depolymerization in a nucleotide-specific fashion^17, 18^. While twinfilin accelerates depolymerization of newly assembled (ADP-P_i_) filaments, it slows down depolymerization of aged (ADP) actin filaments^18^. Notably, twinfilin’s depolymerization activities persist even in cytosol-mimicking conditions i.e., presence of high concentrations of actin monomers^17, 18, 20^.

Although the individual effects of formin, CP and twinfilin on actin assembly are relatively well-characterized, how they simultaneously act in multiprotein teams at barbed ends to influence actin assembly is still poorly understood. This question is especially important as these factors are often found in the same cellular compartments such as filopodia, lamellipodia and stereocilia^6, 18, 21–24^. In addition to twinfilin’s monomer sequestration and barbed end depolymerization functions, it can directly bind CP via its C-terminal tail and accelerate CP’s removal from barbed ends, ultimately reducing CP’s barbed-end dwell time by about 6-fold^20, 25^. The CP-twinfilin interaction has been demonstrated to be required for proper intracellular localization of CP but is dispensable for twinfilin’s uncapping *in vitro* ^20, 26^. Moreover, in absence of direct visualization of CP’s uncapping by twinfilin, the underlying mechanism remains unclear. While a number of recent studies have looked at this twinfilin-CP interaction, twinfilin’s effects on formin have not been extensively studied. We previously reported that formin’s rate of barbed end elongation is not affected by high concentration of twinfilin^18^. Nevertheless, twinfilin’s effects on formin’s long-lived residence at the barbed end has yet to be investigated.

CP and formin were initially thought to bind barbed ends in a mutually exclusive fashion^27–29^. Contrary to this assumption, it was discovered that formin and CP simultaneously bind the same barbed end to form a so-called barbed-end “decision complex” ^30, 31^. Their concurrent presence at the filament end accelerates barbed-end dissociation of formin and CP by 50-fold and 10-fold, respectively^30^. While the mechanisms discussed above reduce barbed-end residence times of formin and CP from 120 minutes and 30 minutes to just a few minutes^30^, these time-scales are still too slow to explain the rapid rates at which intracellular actin structures get assembled, arrested and turned over in a few seconds^1, 32^. This prompted us to take a fresh look at the multicomponent dynamics between CP, formin and twinfilin at filament barbed ends.

Here we show that while twinfilin alone has no direct effects on formin’s processivity, it greatly enhances formin’s processivity in presence of CP (Fig. 1). Using microfluidics-assisted total internal reflection fluorescence (mf-TIRF) microscopy^33, 34^, we find that despite its depolymerization effects on free barbed ends, twinfilin effectively promotes actin assembly when both formin and CP are present together (Fig 2). We find that formin, CP and twinfilin can all simultaneously bind a filament barbed end in a tripartite complex (Fig. 3). The dynamics of this formin-CP-twinfilin complex at the barbed end (BFCT) was visualized by multicolor single molecule imaging. We discovered that twinfilin reduces the lifetime of the CP-formin decision complex at the barbed end by about 17-fold. Our is the first report of a polymerase, a depolymerase and a capper simultaneously occupying the same barbed end. This trimeric complex leads to accelerated transitions between these regulatory proteins at the barbed end. We then directly visualized twinfilin’s uncapping of CP-bound barbed ends and found that twinfilin displaces CP from formin-CP complex much more efficiently than from barbed ends bound to CP alone (Fig 4). While only a single twinfilin binding event is sufficient to remove CP from formin-CP complexes, on average it takes about 30 twinfilin binding events to uncap CP-capped barbed ends. Using separation-of-function mutants, we found that twinfilin’s direct interaction with the actin filament is critical for its ability to rescue formin’s processivity (Fig. 5). To our knowledge, this is the first ever evidence of a depolymerase, polymerase and a capper simultaneously binding a growing filament end, and the multicomponent mechanism by which they can regulate each other’s activities.

**Fig. 1:**
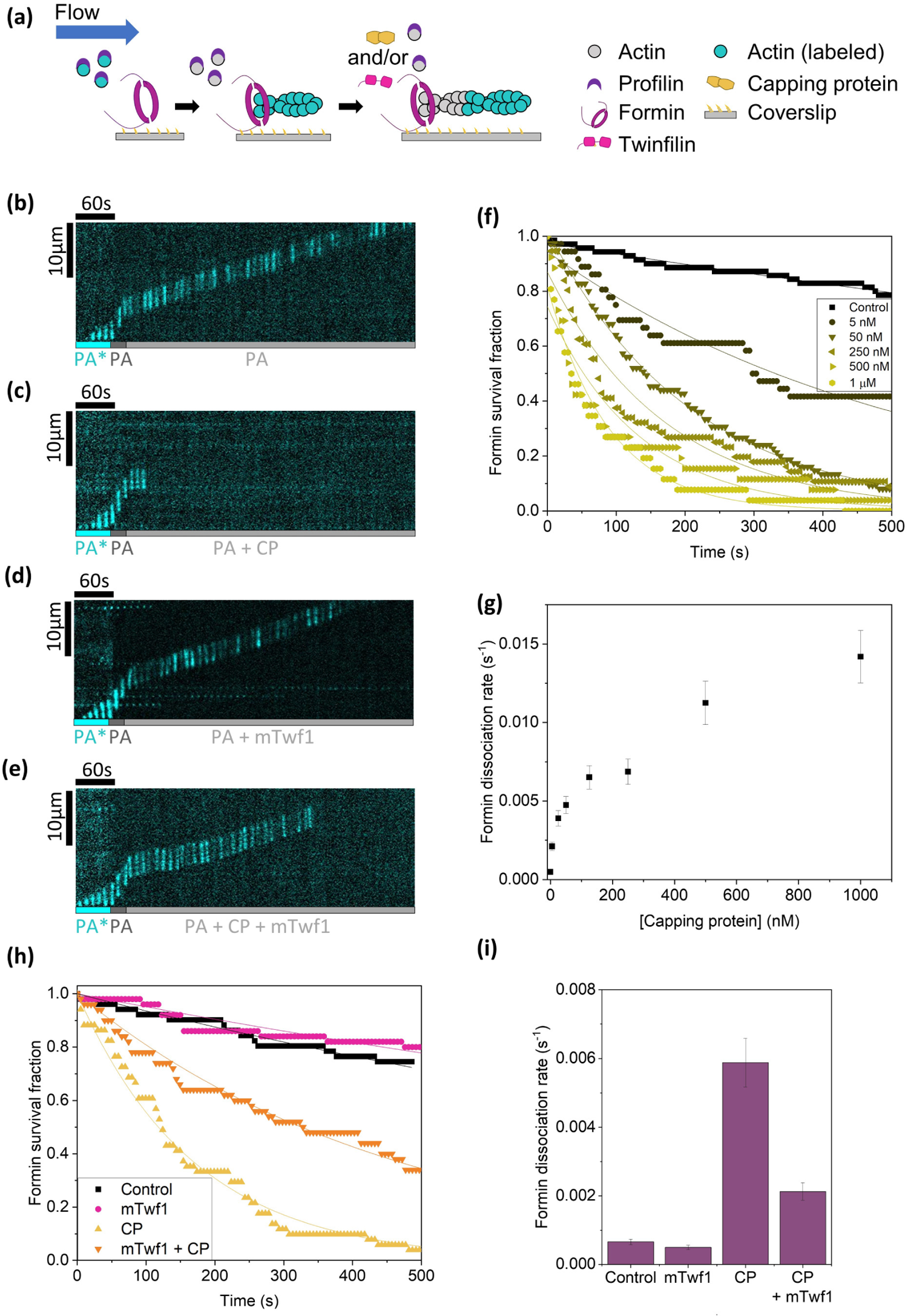
Effect of twinfilin and capping protein (CP) on processivity of formin. **(a)** Schematic representation of the experimental strategy. Actin filaments were nucleated from coverslip-anchored formins by introducing a flow containing 1 µM G-actin (15% Alexa-488 labeled) and 0.5 µM profilin. The filaments were then allowed to elongate in presence of 1 µM unlabeled G-actin and 4 µM profilin to ensure insertional elongation between fluorescent fragment and surface-anchored formins. These filaments were then exposed to a flow containing 0.2 µM unlabeled G-actin, 0.7 µM profilin (control) with or without varying concentrations of CP and/or mTwf1 (alone or together). Survival fraction of filaments attached to formins was monitored as a function of time. **(b)** Representative kymographs of a formin-anchored filament elongating from 0.2 µM unlabeled G-actin and 0.7 µM profilin (PA) (see Supplementary Movie 1). **(c)** Same conditions as (b) but supplemented with 50 nM CP (see Supplementary Movie 2). **(d)** Same conditions as (b) but supplemented with 1 µM mTwf1. **(e)** Same conditions as (b) but supplemented with 50 nM CP and 1 µM mTwf1 (see Supplementary Movie 3). **(f)** Survival fraction of formin-bound filaments (BF) as a function of time in presence of PA supplemented with varying concentrations of CP. Experimental data (symbols) are fitted to a single-exponential decay function (lines) to determine formin dissociation rate *k_-F_* (BF◊B + F). Number of filaments analyzed for each condition (0 to 1 µM CP): 70, 36, 75, 56, 26, 26) **(g)** Formin dissociation rate *k-_F_* as a function of CP concentration, determined from data shown in (f). **(h)** Survival fraction of formin-bound filaments (BF) as a function of time in presence of PA alone (black symbols, n = 51 filaments) or supplemented with 1 µM mTwf1 alone (magenta symbols, n = 50 filaments), 50 nM CP alone (yellow symbols, n = 51 filaments) or both 1 µM mTwf1 and 50 nM CP (orange symbols, n = 50 filaments). Experimental data (symbols) are fitted to a single-exponential decay function (lines). **(i)** Formin dissociation rate *k_-F_* as determined from data in (h). Error bars indicate 65% confidence intervals based on fits (see methods).

**Fig. 2:**
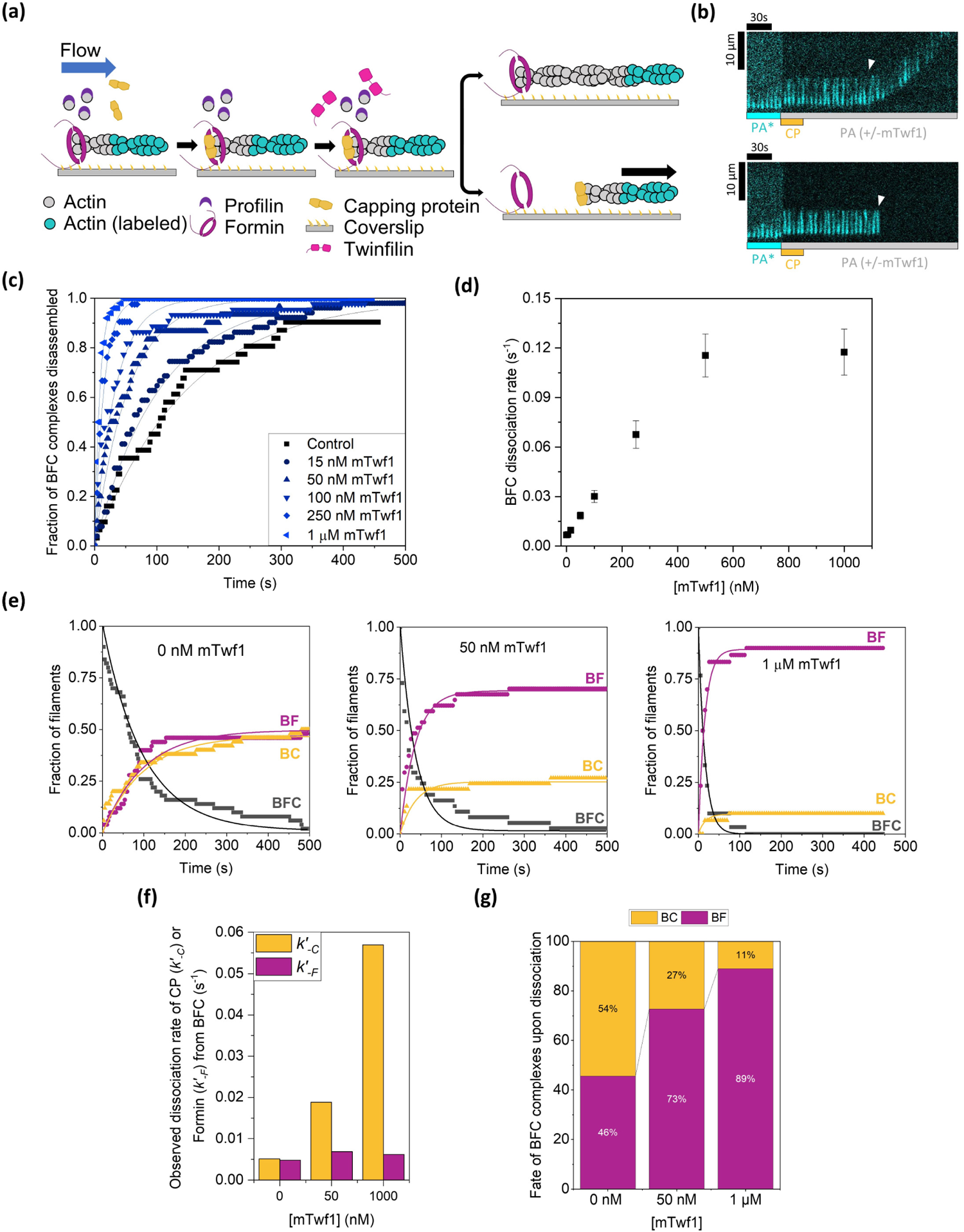
Effect of twinfilin on capping protein (CP) – formin “decision complex”. **(a)** Schematic representation of the experimental strategy. Actin filaments were nucleated from coverslip-anchored formins by introducing a flow containing 1 µM G-actin (15% Alexa-488 labeled) and 0.5 µM profilin. The filaments were then exposed to a flow containing 1 µM unlabeled G-actin, 4 µM profilin and 1 µM CP for about 10 s to convert formin-bound barbed ends (BF) to formin-CP bound barbed ends or “decision complexes” (BF + C ◊BFC). These BFC complexes were then exposed to a flow containing PA only or supplemented with varying concentrations of mTwf1. **(b)** Representative kymographs of two formin-elongating filaments transitioning to paused BFC state upon exposure to 1 µM CP. Upon removal of CP and exposure to PA (with or without mTwf1), filaments either resume elongation due to dissociation of CP from BFC (top, BFC ◊ BF + C) or detach (bottom, BFC ◊ BC + F). White arrowheads denote the moment formin or CP dissociation from the BFC complex. **(c)** Survival fraction of formin-CP bound filaments (BFC) as a function of time in presence of PA with/without varying concentrations of mTwf1. Experimental data (symbols) are fitted to a single-exponential function (lines) to determine the rate of BFC disassembling into BF or BC. Number of filaments analyzed for each condition (0 to 1 µM mTwf1): 31, 51, 30, 44, 42, 50). **(d)** Rate of BFC dissociation into BC or BF as a function of mTwf1 concentration, determined from data in (c). **(e)** Fraction of BFC (black symbols) filaments transitioning either to BF (filaments resuming elongation, magenta symbols) or BC (filaments detached, yellow symbols). The experimental data (symbols) are fitted to exponential fits (lines), such that *k_-BFC_* = *k’_-F_* + *k’_-C_*, where *k’_-F_* is the dissociation rate of formin from BFC (BFC ◊ BC + F) and *k’_-C_* is the dissociation rate of CP from BFC (BFC ◊ BF + C). Conditions – left (0 nM mTwf1, 50 filaments), center (50 nM mTwf1, 37 filaments), right (1 µM mTwf1, 30 filaments). See Supplementary Fig. 2 for the entire range of mTwf1 concentrations. **(f)** Dissociation rate of CP (*k’_-C_*) or Formin (*k’_-F_*) from BFC decision complexes **(g)** Percentages of BFC complexes shown in (e) transitioning to BC (yellow) or BF (magenta) respectively at different mTwf1 concentrations. Error bars indicate 65% confidence intervals based on fits (see methods).

**Fig. 3:**
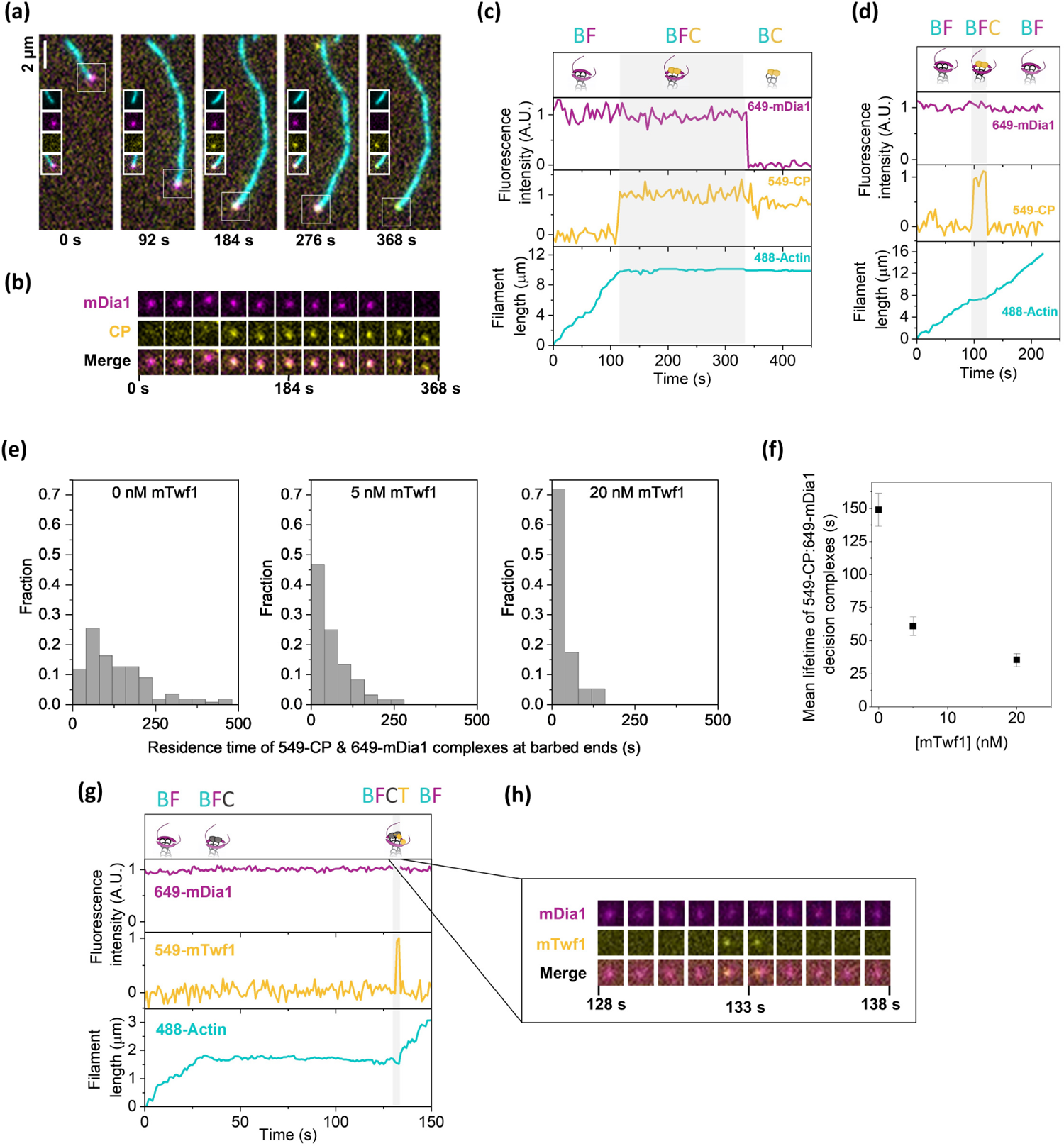
Direct visualization of formin, CP and mTwf1 at barbed ends. **(a)** Representative time-lapse images of a multi-color single-molecule TIRF experiment (see Supplementary Movie 4). Actin filaments were initiated by incubation of 0.5 µM G-actin (15% Alexa-488 labeled, 0.5% biotin labeled), 1 µM profilin and 50 pM 649-mDia1 (magenta). These 649-mDia1 bound filaments were then exposed to PA, 10 nM 549-CP (yellow) with or without mTwf1. The white box around the filament end in each frame indicates the location of the barbed end. Insets show individual and merged channels localized at the barbed end. Scale bar, 2 µm. **(b)** Fluorescence images of a 13 x 13-pixel box around the barbed end of the filament from (a) showing formation and dissociation of a mDia1– CP complex at the barbed end. Formin channel is shown in magenta and CP channel is shown in yellow. **(c)** Fluorescence intensity and filament length records of the filament in (a) with formation and resolution of the “decision complex” in absence of mTwf1. **(d)** Same as c but in presence of 20 nM mTwf1. The gray shaded boxes indicate the period when both 549-CP and 649-mDia1 were simultaneously present at the barbed end i.e., BFC complex was intact. **(e)** Distribution of lifetimes of 649-mDia1:549-CP decision complexes at barbed ends in presence of 0 nM mTwf1 (left, n= 110 BFC complexes), 5 nM mTwf1 (center, n= 60 BFC complexes) and 20 nM mTwf1 (right, n= 57 BFC complexes). **(f)** Mean lifetimes (± sem) of 649-mDia1:549-CP decision complexes at barbed ends as a function of mTwf1 concentration, determined from data in (e). **(g)** Fluorescence intensity and filament length records of a filament with formation and resolution of the “decision complex” formed in presence 100 pM 649-mDia1, 10 nM unlabeled CP and 40 nM 549-mTwf1. The gray shaded box indicates the time duration when both 549-mTwf1, 649-mDia1 and unlabeled CP were simultaneously present at the barbed end. **(h)** Cropped fluorescence images of a 13 x 13-pixel box around the barbed end of the filament in (g) showing formation and dissociation of a 649-mDia1– unlabeled CP complex in presence of 549-mTwf1. The formin channel is shown in magenta and twinfilin channel is shown in yellow.

**Fig. 4:**
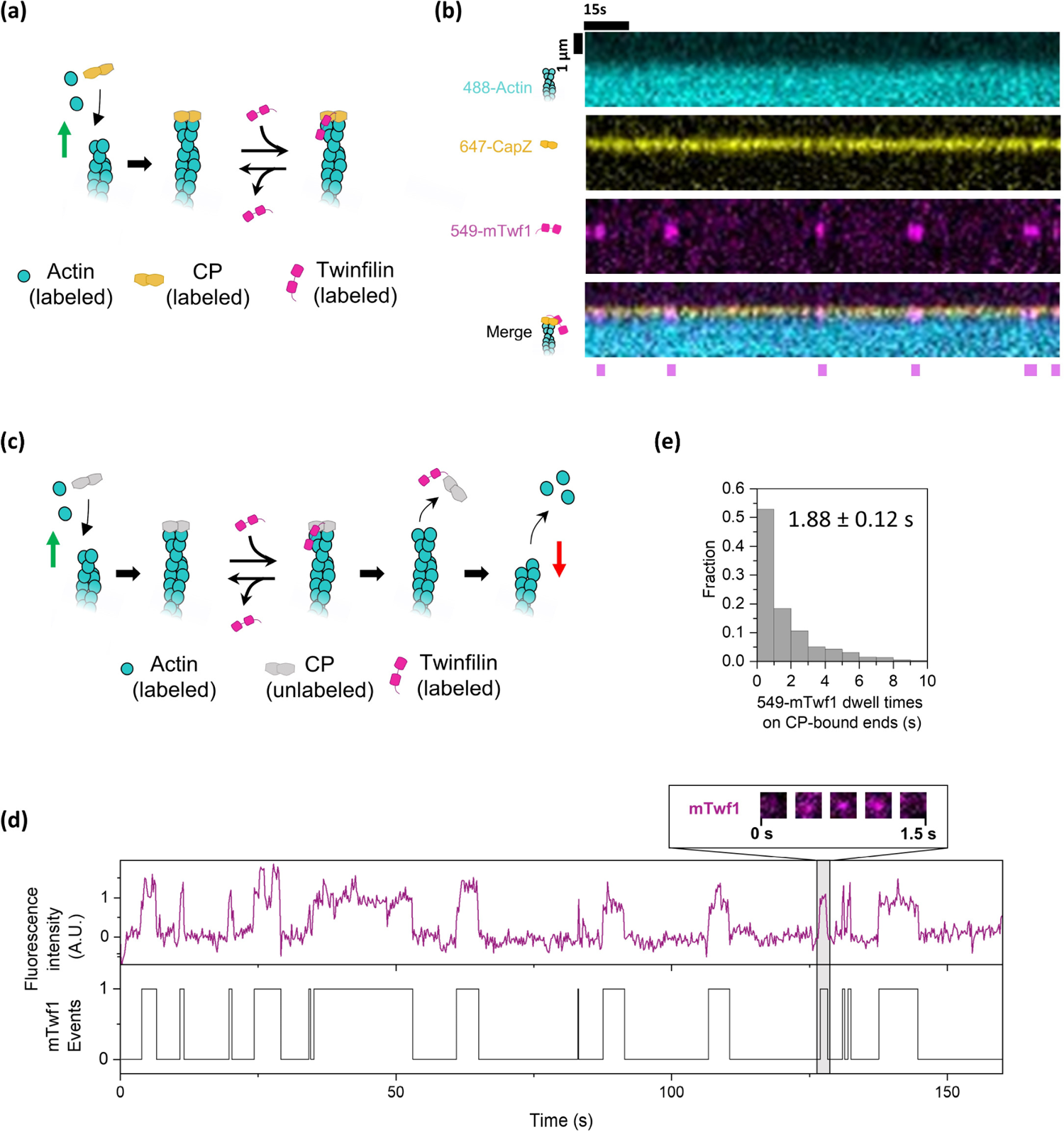
Visualization and characterization of twinfilin on CP-bound barbed ends. **(a)** Schematic of the single-molecule experiment with labeled CP and labeled mTwf1. Actin filaments were assembled from of 1 µM G-actin (15% Alexa-488 labeled, 1.4% biotin labeled) and sequentially capped by 10 nM 649-CP. These filaments were then exposed to 20 nM 549-mTwf1. Green arrow denotes polymerization. **(b)** Binding of labeled twinfilin on CP-bound filament barbed ends recorded at 1 s time resolution. Kymographs show Alexa-488 actin (top), 649-CP (second from top), 549-mTwf1 (third from top) and merge (bottom). Magenta bars denote episodes in which a 549-mTwf1 molecule was present at the 649-CP bound barbed end of the filament. **(c)** Schematic of the experimental set up. Actin filaments were polymerized as in (a), and capped by unlabeled CP. These filaments were then exposed to 15 nM 549-mTwf1. Arrival and departure of 549-mTwf1 molecules at the barbed end were recorded until the filament started depolymerizing i.e., CP dissociated. Green arrow denotes polymerization, and the red arrow denotes depolymerization. **(d)** Distribution of residence times 549-mTwf1 on unlabeled CP-bound filament barbed ends (n= 510 binding events across 16 filaments). **(e)** Top: time records of 549-mTwf1 fluorescence intensity at the unlabeled CP-bound barbed end of an actin filament. Intensity is integrated over a 5 × 5 pixel square centered around the barbed end of the filament. Bottom: Presence (1) or absence (0) of 549-mTwf1 molecules at the barbed end of the filament shown above. Inset: cropped fluorescence images of a 10 x 10-pixel box around the barbed end of the filament showing the arrival, presence and departure of a single 549-mTwf1 molecule at CP-bound barbed end.

**Fig. 5:**
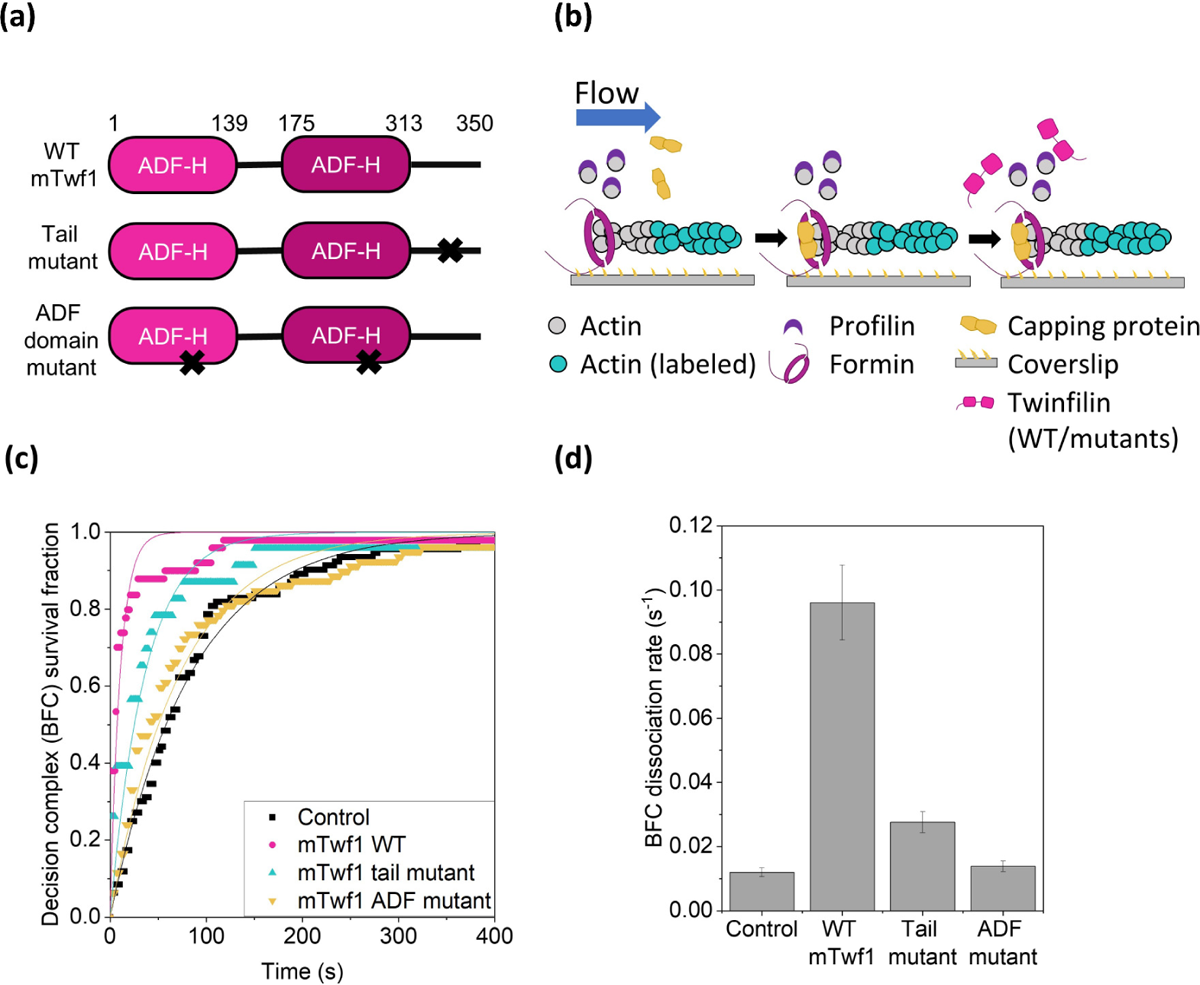
Twinfilin’s direct interaction with actin filament is essential for its effects on formin-CP decision complexes. **(a)** Domain diagram of wild-type and mutant mTwf1 constructs used here (diagram adapted from^20^). “x” denotes the location of mutations **(b)** Schematic representation of the experimental strategy. Actin filaments were nucleated from coverslip-anchored formins by introducing a flow containing 1 µM G-actin (15% Alexa-488 labeled) and 0.5 µM profilin. The filaments were then exposed to a flow containing 1 µM unlabeled G-actin, 4 µM profilin and 500 nM CP for about 10 s to convert formin-bound barbed ends (BF) to formin-CP bound barbed ends or “decision complexes” (BF + C ◊BFC). These BFC complexes were then exposed to a flow containing PA only or supplemented with wild type or mutant mTwf1. **(c)** Survival fraction of formin-CP bound filaments (BFC complexes) as a function of time in presence of PA only (black symbols, 92 filaments), or supplemented with 1 µM wild type mTwf1 (black, 36 filaments), mTwf1 ADF mutant (green, 77 filaments) or mTwf1 tail mutant (blue symbols, 22 filaments). Experimental data (symbols) are fitted to a single-exponential function (lines) to determine BFC dissociation rate *k_-BFC_*. Error bars indicate 65% confidence intervals based on fits (see methods). **(d)** BFC dissociation rate for wild type and mutant mTwf1, determined from data in (c). Error bars indicate 65% confidence intervals based on fits (see methods).

## Results

### Capping protein and twinfilin differentially influence formin’s processivity

We studied how CP and twinfilin influence formin’s processivity at actin filament barbed ends. Fluorescent actin filaments were initiated in a mf-TIRF chamber by exposing coverslip-anchored formins (mDia1 FH1-FH2-C) to a solution containing fluorescent actin monomers and profilin-actin (PA) (Fig. 1a). Then, to confirm these filaments were indeed elongating from formins on the glass coverslip, a solution containing profilin and unlabeled actin monomers was flowed. Since elongation occurs by insertion of unlabeled monomers at the formin bound barbed end, the pre-existing fluorescent segment of the filament appears to move in the direction of the flow (Fig. 1a, b; Supplementary movie 1), away from the location of formin anchoring.

Dissociation of surface-anchored formins from barbed ends caused immediate disappearance of filaments from the field of view (BF → B + F, where B denotes the barbed end, F denotes the formin and BF is the formin-bound barbed end).

We recorded the disappearance of actin filaments in the field of view over time to quantify formin’s processivity. Changes in survival fraction of formin-bound filaments were then used to determine the observed dissociation rate of formin from the barbed end. Using unlabeled actin subunits eliminates the effects of labeled actin on formin’s processivity. To confirm that the filament disappearance was due to detachment of filaments from the formin rather than due to detachment of the entire formin-filament complex from the glass coverslip, we re-exposed the surface to a flow containing 1 µM Alexa-488 G-actin and 0.5 µM profilin. Consistent with previous studies, over 80% of formins were able to renucleate new filaments following detachment of initially nucleated filaments^30, 35^.

First, we asked how CP influences processivity of formins on filament barbed ends. Formin-elongated fluorescent filaments were exposed to a solution containing either profilin-actin alone, or with varying concentrations of CP. In control reactions, the fluorescent segment of actin filaments continued to move away from the attached formin, at a constant speed, in the direction of the flow, indicating processive elongation by formin (Fig. 1b, Supplementary movie 1). Only a small fraction (∼20 %) of filaments dissociated from formins during the duration of the experiment (500 s). The average barbed-end dwell time of formin mDia1 was ∼35 min in control experiments (Fig 1f, dwell time = 1/(dissociation rate)). Formin’s long barbed-end residence time measured here agrees with previous studies^13, 14, 35^. Interestingly, when capping protein was introduced (in presence of PA), actin filaments rapidly dissociated from formins and disappeared from the field of view (Fig. 1c, f; Supplementary movie 2). The rate of dissociation of formins increased with CP concentration (5 nM to 1 µM). Compared to control, 1 µM CP increased formin’s rate of dissociation from the barbed end by about 30-fold (Fig. 1g).

While the cytoplasmic concentration of CP is around 1 µM^36^, the majority of it is thought to be sequestrated by V1/myotrophin^37^. As a result, only about 10 – 20 nM free CP is expected to be available for binding barbed ends in cells ^37^. We found that even at low CP concentrations in this range (∼50 nM), CP was still able to accelerate formin dissociation by about 10-fold compared to the control (Fig. 1g). This means that even low concentrations of active CP were sufficient to rapidly terminate formin-induced actin assembly as compared to control.

We then asked if twinfilin might also influence formin’s association with barbed ends. In contrast to capping protein, we found that presence of up to 1 µM twinfilin (mouse mTwf1, referred to as ‘twinfilin’ hereafter) did not significantly change formin’s barbed-end dwell time (Fig. 1d, h, i). Consistent with our earlier study, twinfilin also had no observable effect on formin’s rate of elongation^18^ (Supplementary Fig. 1). Our results imply that unlike capping protein, formins fully protect barbed ends from twinfilin, indicating that twinfilin cannot associate with a formin-bound barbed end.

So far, we have explored the effects of CP and twinfilin individually on formin. CP and twinfilin are, however, simultaneously present in the same cellular compartments, such as filopodia, lamellipodia etc., which also contain formin-elongating actin filaments. We therefore asked how simultaneous presence of twinfilin and CP influences formin’s processivity. When formin-nucleated filaments are exposed to CP and twinfilin at the same time (in presence of PA), formin dissociates at a rate intermediate between that of control and capping protein (Fig. 1h, i; Supplementary movie 3). While there was a 10-fold reduction in formin’s processivity in presence of 50 nM CP, the reduction was just 3-fold when 50 nM CP was supplemented with 1 µM twinfilin. Our data suggests that twinfilin’s presence led to a reduction in the adverse effects of CP on formin’s processivity. Consistently, actin filaments also grew substantially longer (Fig. 1e) prior to their detachment in presence of both twinfilin and CP as compared to CP alone (Fig. 1c).

### Twinfilin accelerates resolution of formin-CP BFC complex

X-ray diffraction and EM studies suggest that both CP and twinfilin interact with the last two subunits at the filament barbed end^25, 38^. Moreover, twinfilin destabilizes CP’s barbed end localization leading to a 6-fold reduction in CP’s barbed-end lifetime^20^. We therefore wondered if twinfilin’s rescue of formin’s processivity from CP might be due to its destabilizing effects on CP in the formin-CP-barbed end “decision complex.” To investigate this, we formed decision complexes by exposing formin-nucleated fluorescent actin filaments to a high concentration of CP (1 µM). This caused an almost immediate arrest of filament elongation and formation of formin-CP-barbed end decision complexes^30, 31^ (referred to as BFC henceforth, where B is the barbed end, F is formin and C is CP). To study the effects of twinfilin on BFC complexes, BFC filaments were exposed to a flow containing either profilin-actin only (control) or supplemented with varying concentrations of twinfilin (Fig. 2a).

The BFC complexes can dissociate by two different routes – 1) via dissociation of CP (BFC → BF + C, with CP departing from BFC with a rate *k’_-C_*), which leads to resumption of elongation of formin-bound filaments (Fig. 2b, top) or 2) via dissociation of formin (BFC → BC + F, with formin detaching from BFC with a rate *k’_-F_*), which leads to detachment and disappearance of the filament (Fig. 2b, bottom). We found that twinfilin dramatically accelerated the disassembly of BFC complexes in a concentration-dependent manner (Fig. 2c, d). Compared to control, 1 µM twinfilin increased the rate of BFC dissociation (*k_-BFC_*) by about 18-fold.

The dissociation kinetics of BFC complexes into BC and BF were exponential with observed rate *k_-BFC_* = *k’_-F_ + k’_-C_* where *k’_-F_* is the observed rate of dissociation of formin from the BFC complex and *k’_-C_* is the observed rate of dissociation of CP from the BFC complex (Fig. 2e, Supplementary Fig. 2). The fraction of filaments transitioning to BF and BC states upon BFC dissociation is given by the *k’_-C_/(k’_-C_ + k’_-F_)* and *k’_-F_/ (k’_-C_ + k’_-F_)* (see methods). We derived the values *k’_-C_* and *k’_-F_* as a function of twinfilin concentration (Fig. 2e, f, Supplementary Fig. 2) and found that while 1 µM twinfilin increased CP’s dissociation rate from the BFC complex by ∼11-fold, formin’s rate of dissociation did not change significantly (Fig. 2f). We also analyzed the route by which the BFC complexes dissociated (BFC → BF + C or BFC → BC + F). In absence of twinfilin, slightly more than half of all BFC complexes transitioned to BC (∼54%) and the rest transitioned to BF (∼46%). In presence of twinfilin however, this ratio was skewed heavily towards BF (∼90%) i.e., CP dissociated and left formin solely bound to the barbed end (Fig. 2g). These filaments immediately returned to rapid elongation, characteristic of formin’s presence at the barbed end.

Taken together, our results indicate that although twinfilin is a depolymerase of free barbed ends, it promotes filament assembly by formin when both CP and formin are present. Just as twinfilin can uncap CP from free barbed ends^20^, our results indicate twinfilin might also be able to destabilize CP from BFC complexes by forming a ternary complex, BFCT, with formin and CP at the barbed end (T in BFCT denotes twinfilin). However due to our inability to directly visualize twinfilin, it was not possible to ascertain whether twinfilin’s effects were due to its barbed end binding or its interactions with the filament side.

### Single-molecule visualization of twinfilin’s effects on BFC dynamics

To directly visualize the effects of twinfilin on dynamics of the BFC complex we used three-color single molecule TIRF imaging. SNAP-tagged constructs of CP (SNAP-CP) and formin (SNAP-mDia1) were expressed and labeled with benzylguanine functionalized green-excitable (549-CP) and red-excitable (649-mDia1) fluorescent dyes. Photobleaching data confirmed that majority of 549-CP molecules were labelled with only one dye molecule, consistent with a single SNAP tag per heterodimeric CP molecule ^31^ (Supplementary Fig. 3). Photobleaching tests of 649-mDia1 showed that majority of these molecules exhibited a single- or double-step bleaching profile, consistent with the dimeric nature of formin mDia1 molecules ^30, 39^ (Supplementary Fig. 4). Alexa-488 labeled G-actin actin monomers were incubated with 649-mDia1 in a non-microfluidic, conventional flow cell. The majority of newly nucleated filaments displayed fluorescent formins at their barbed ends (Fig. 3a). All filament barbed ends with detectable 649-mDia1 underwent rapid, continuous elongation with no noticeable pauses. Consistent with long barbed-end dwell times and low photobleaching, the majority of 649-mDia1 molecules remained bound to elongating filaments for the duration of the experiment.

649-mDia1 bound barbed ends were exposed to a profilin-actin solution containing either 549-CP alone or together with twinfilin. In absence of twinfilin, we routinely observed the following (Fig. 3a, b) – first, the 649-mDia1 was joined at the barbed end by a 549-CP molecule to form a BFC complex; second, upon arrival of 549-CP the filament immediately stopped elongating; third, following a brief interval of time during which both the molecules were jointly bound to the barbed end, the 649-mDia1:549-CP complex dissociated and one of these two molecules departed from the barbed end, leaving the other behind (Supplementary movie 4). In approximately 58% of the complexes 649-mDia1 dissociated (N = 42 out of 72 complexes), leaving 549-CP at the barbed end, and no further elongation was observed (Fig. 3a, c). In the remaining complexes, 549-CP departed (N=30 out of 72 complexes) and the filament immediately switched from the paused state to the rapidly elongating state, characteristic of formin-bound barbed ends. The 649-mDia1:549-CP barbed end complex had an average lifetime of 149 ± 12.4 s (Figure 3e, left). In contrast, when 649-mDia1 bound filaments were exposed to 549-CP but in presence of 20 nM twinfilin, a much shorter pause was observed during which both 549-CP and 649-mDia1 were bound and the filament elongation was arrested (Fig. 3d). The average complex lifetime reduced to 35.4 ± 5 s, a 75% reduction over control (Fig. 3e, f). In agreement with our mf-TIRF experiments, twinfilin also altered the outcome of decision complex resolution. About 73% of complexes (36 out of 49) transitioned to the 649-mDia1 BF state with 20 nM twinfilin as compared to 42% of complexes (30 out of 72) in control experiments.

The mf-TIRF and single-molecule experiments together conclusively establish that twinfilin influences both the lifetime and eventual outcome (BF or BC) of the BFC decision complex dissociation. We then asked if these effects are caused by binding of twinfilin along the filament length or by its interactions at the barbed end. To do this, we expressed mouse twinfilin-1 as a SNAP-tagged fusion protein and directly visualized its interactions with BFC complex (see methods). SNAP-tagging didn’t alter mTwf1’s ability to uncap CP from CP-bound barbed ends (Supplementary Fig. 5). Photobleaching records of surface adsorbed 549-SNAP-mTwf1 showed that almost all molecules exhibited a single-step photobleaching, confirming that labeled twinfilin molecules were monomeric (Supplementary Fig. 6).

Using labeled twinfilin, we further examined the mechanism by which twinfilin accelerated dissociation of the BFC complex. We exposed 649-mDia1 bound Alexa-488 actin filaments to a solution containing fluorescently labeled 549-mTwf1 and unlabeled CP. Expectedly, filaments rapidly paused due to decision complex formation between unlabeled CP and 649-mDia1, leading to an abrupt stop in translocation of 649-mDia1 molecules bound to actin filaments. To our surprise, we observed 549-mTwf1 molecules were now able to transiently join 649-mDia1 (and unlabeled CP) at the barbed end (Fig. 3g, h). Association of 549-mTwf1 molecules with 649-mDia1:unlabeled CP bound barbed ends was very brief, with an average dwell time of 1.4±0.2 s (n = 8 events), translating to a dissociation rate of 0.71 s^−1^. The majority of 549-mTwf1:649-mDia1:unlabeled CP BFCT complexes remained intact just for a single frame implying the 0.71 s^-1^ rate of dissociation of 549-mTwf1 molecules from BFCT complex is the lower bound for this rate. Importantly, the departure of 549-mTwf1 from the decision complex led to an immediate transition of the barbed end from the arrested state to the fast formin-based elongation and translocation of the 649-mDia1 molecule. Notably, in absence of CP, we never observed colocalization of 649-mDia1 and 549-mTwf1 at barbed ends. This explains our mf-TIRF results that twinfilin doesn’t influence the rate of elongation or processivity of formins (Fig. 1h, Supplementary Fig. 1). Together, these observations indicate that the unlabeled CP molecule and twinfilin departed the barbed end simultaneously, leaving formin behind. Notably, we never observed twinfilin and CP simultaneously arrive at formin-occupied barbed ends.

### Single molecule analysis uncovers twinfilin’s uncapping mechanism

Although twinfilin is known to accelerate CP’s dissociation ^40^, in absence of simultaneous visualization of these proteins at barbed ends, the underlying mechanism has remained obscure. We directly visualized simultaneous association of CP and twinfilin at barbed ends (Fig 4a). Actin filaments assembled from Alexa-488 labeled actin monomers were transiently exposed 649-CP in a non-microfluidic, conventional flow cell. All filaments with a visible 649-CP signal at their barbed end immediately stopped growing following CP’s arrival. Upon exposure to 549-mTwf1, short-lived associations of twinfilin at the 649-CP-bound barbed ends were seen. To our surprise, unlike in the case of BFC complexes where a single twinfilin binding event was sufficient to cause CP’s departure, 649-CP remained bound to the barbed end despite repeated barbed end arrivals and departures of 549-mTwf1 molecules (Fig. 4b). Each 549-mTwf1:649-CP colocalization event lasted about a second. However, owing to slow rate of uncapping by twinfilin (it only accelerates uncapping rate by 6-fold) and transient nature of these interactions, watching simultaneous departure of 649-CP and 549-mTwf1 required frequent observation (several frames per second) for long durations. This excessive exposure to light led to the disappearance of the 649-CP signal from barbed ends and in many cases without the filament beginning to depolymerize. This suggests that the disappearance of the 649-CP signal was due to photobleaching and not due CP dissociation. We therefore decided to employ an alternative strategy wherein filaments capped with unlabeled CP were exposed to 549-mTwf1 (Fig. 4c). Importantly, during exposure to 549-mTwf1, no free CP or actin monomers were present in the chamber. Barbed end depolymerization of filaments following CP’s dissociation was used as a readout for CP’s departure from the barbed end (Fig. 4c). We found that on average it took about 30.9 ± 6.5 successive 549-mTwf1 binding events at the barbed end before CP’s dissociation (as determined by beginning of barbed end depolymerization) (Fig. 4d). Upon binding, a 549-mTwf1 molecule remained bound for about 1.9 ± 0.1s (n = 510 events) before dissociating (Fig. 4e). Notably, 549-mTwf1 intensity appeared and disappeared from the barbed end in a single step, suggesting only a single 549-mTwf1 molecule was able to colocalize with a CP molecule at the barbed end (Fig. 4d).

### Both twinfilin’s CP-binding and actin-binding domains are necessary for its effects on BFC dynamics

As mentioned earlier, twinfilin can directly bind CP as well as interact with terminal F-actin subunits at the barbed end^25, 41^. Which of these interactions is responsible for twinfilin’s effects on BFC complex dynamics? To answer this question, we purified two twinfilin mutants: 1) the “ADF domain mutant”, which inhibits twinfilin’s interaction with actin^42^ and 2) the “tail mutant”, which interferes with twinfilin’s direct binding to CP (Fig. 5a). Pre-formed BFC decision complexes (similar to the strategy used in Fig. 2a) were exposed to a solution containing either profilin-actin only or supplemented with wild-type or mutant twinfilin (see schematic in Fig. 5b). Mutations in the twinfilin tail led to BFC dissociation activity partway between control and the wild-type (Fig. 5c, d). However, the ADF-domain mutant did not accelerate BFC complex dissociation, implying that direct contact between twinfilin and actin is essential for these activities. Consistently, an earlier study showed that the ADF-domain mutant is also incapable of uncapping CP from free barbed ends^20^. Taken together, our data suggest that while twinfilin’s interaction with CP via its tail domain is beneficial, twinfilin binding to actin via its ADF domains is necessary for it to be able to form the BFCT ternary complex at the barbed end and for its ability to rapidly dissociate BFC decision complexes.

## Discussion

Living cells can rapidly tune assembly of their actin networks in response to external stimuli^1^. The actin filament barbed end is considered to be the primary site of regulation of actin assembly. Over the last few decades, several barbed-end interacting factors like polymerases (e.g., formin), cappers (e.g., CP) and depolymerases (e.g., twinfilin) have been identified and characterized. Alone, formin enhances the rate of filament assembly by up to 5-fold^13^, CP arrests growth ^16^ and twinfilin promotes depolymerization^17, 18^. How these distinct protein activities get integrated into the complex, intracellular milieu of a shared cytoplasm remains an open question. This is especially important in cellular compartments like stereocilia, filopodia, and lamellipodia^6, 18, 20–24^ where these proteins co-exist. We have uncovered a new mechanism by which twinfilin, formin and CP simultaneously bind a filament barbed end to form a novel multicomponent protein ecosystem.

We find that CP on its own increases the rate of dissociation of formin from barbed ends, leaving behind CP-capped filaments unable to grow (Figure 1c and 3a, c). On the other hand, twinfilin depolymerizes free barbed ends even in the presence of polymerizable actin monomers^17, 18, 20^. In such conditions, where capping and disassembly are strongly favored over assembly, how would actin assembly occur at all? Combining our observations with previous studies, we present a schematic for how polymerases, depolymerases and cappers simultaneously regulate actin assembly (Fig. 6). Filament assembly outcomes would depend upon whether the filament was elongating freely or bound to a formin. Free barbed ends would rapidly get capped by CP (Fig. 6a). Twinfilin would then catalyze CP’s dissociation from the end, leaving twinfilin alone at the depolymerizing barbed end^20^. Formin bound barbed ends on the other hand would get paused by CP to form the BFC decision complex (Fig. 6b)^30, 31^. Twinfilin’s binding to BFC to form BFCT complex would cause CP’s dissociation and resumption of formin-based filament elongation. As the filament ages, other actin binding proteins with preference for ADP-F-actin e.g., cofilin and cyclase associated protein (CAP), would initiate severing and pointed-end depolymerization of the filament^43–46^. As a result, while filaments with free barbed ends would rapidly disassemble into monomers due to depolymerization from both ends, formin bound ends in presence of twinfilin would persistently grow at their barbed ends and depolymerize from their pointed ends, akin to treadmilling. In summary, our observations suggest that the depolymerase, twinfilin, promotes assembly of formin-bound filaments over filaments with unprotected free barbed ends.

**Fig. 6:**
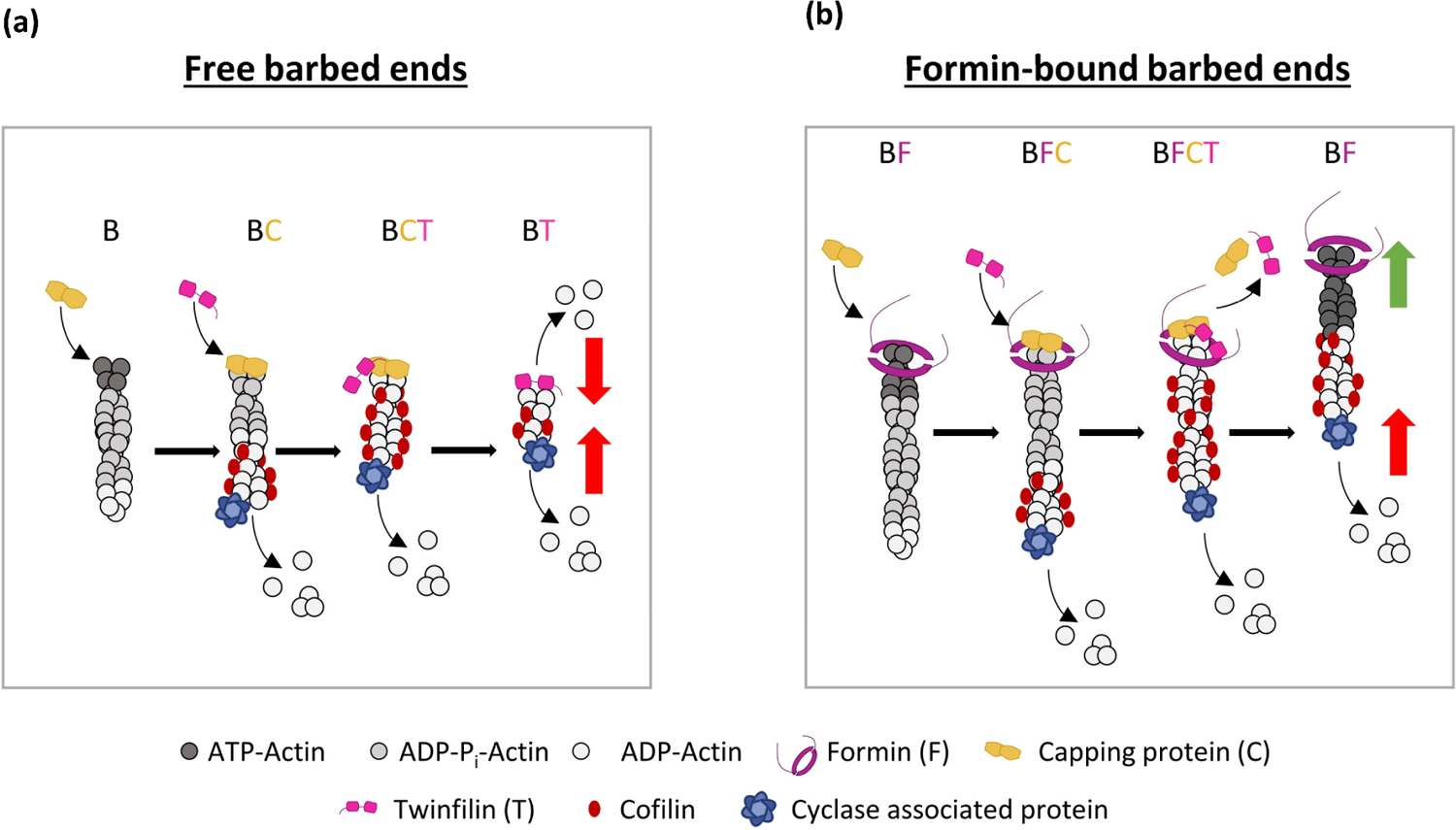
Schematic for twinfilin, formin and CP’s simultaneous regulation of barbed end dynamics. Barbed end outcomes would differ depending upon the presence or absence of formin at the barbed ends. **(a)** Free barbed ends (B) would rapidly get capped by CP (C), followed by CP’s dissociation by twinfilin (T). This would leave twinfilin alone at the filament end resulting in barbed end depolymerization of the filament. At the same time, cofilin would bind the sides of the aging filament and synergize with cyclase associated protein (CAP) to initiate pointed-end depolymerization of the filament. The simultaneous depolymerization at the two ends would result in complete disassembly of the filaments into monomers. **(b)** Formin (F) bound barbed ends would get paused by CP to form the decision complex. Twinfilin’s binding to the decision complexes would cause CP’s dissociation and renewal of formin-based filament elongation. As a result, the filament would appear to treadmill i.e., continue elongating at the barbed end while at the same time being depolymerized at the pointed end by CAP-cofilin synergy. “B” denotes barbed end, “F” denotes formin, “C” denotes CP and “T” denotes twinfilin.

In addition to being a depolymerase, twinfilin is also a barbed end uncapper. Our single-molecule experiments bring new insights into twinfilin’s uncapping mechanism. We found that unlike cofilin, which uncaps by decorating actin filament sides^46^, twinfilin directly associates with CP-bound barbed ends to uncap actin filaments. Twinfilin is, at best, a mediocre uncapper on its own as it takes on average 31 twinfilin association and disassociation events at the barbed end per successful uncapping event. In comparison, CARMIL accelerates CP’s departure by about 180-fold, bringing down CP’s barbed-end half-time from 30 min to just about 10 s^47^. As a result, although both CARMIL and twinfilin bind CP similarly via their CPI motifs^48^, they exert vastly different control on CP.

Our earlier study showed that simultaneous presence of CP and formin at the barbed end leads to a weakening of both of their barbed end binding^30^. Our experiments suggest that the already weakly bound CP in BFC complexes gets uncapped by twinfilin much faster than if CP alone was tightly bound to barbed ends. This suggests that formin and twinfilin’s uncapping capabilities could potentially synergize to increase uncapping.

In addition to uncapping, twinfilin is also a known actin depolymerase. How does it carry out both of these activities? Our experiments with twinfilin variants containing mutations in its actin-binding ADF domains suggest that twinfilin’s interactions with the actin filament are essential for its ability to rescue formin’s processivity. Mutations in twinfilin’s tail domain interfere with twinfilin’s binding to CP^20^. Although less strongly than the wild-type, we find that the tail mutant can still dissociate CP from the formin-CP barbed end complex and modestly rescue formin’s processivity. This can be interpreted in two ways. 1) For twinfilin’s effects on the decision complex, its direct binding to CP is beneficial but not necessary. Interestingly, a previous study found that the tail mutant uncaps CP from free barbed ends much faster than the wild type^20^. 2) Alternatively, this could also mean that there are additional sites on twinfilin apart from its tail which help it directly bind CP. The linker region between the two ADF domains in twinfilin has been proposed as an extra site of contact between CP and twinfilin when both are bound to actin^25^. Importantly, mutations in ADF domains, which interfere with twinfilin’s actin binding and extinguish twinfilin’s uncapping activities at free barbed ends^20^, also extinguish twinfilin’s effects on decision complex dissociation. Together, these results indicate that in the ternary BFCT complex comprising of formin, CP and twinfilin at the barbed ends, each factor is directly interacting with the actin filament and not via each other.

Although incredible, twinfilin’s behavior in promoting assembly is not unique. Twinfilin is part of the ADF homology (ADF-H) family of proteins and contains two ADF domains^49^. Like twinfilin, cofilin also promotes uncapping and accelerates filament end depolymerization (on top of its severing activities)^45, 46^. Both of these proteins also synergize with cyclase associated protein (CAP)^17, 43, 44^. It will be interesting to investigate CAP’s effect on decision complex dynamics, especially since it contains WH2 domains which can directly bind actin. A number of other factors including CP, Bud14 and Hof1 inhibit formin-based elongation via diverse mechanisms. CP and Bud14 displace formin from the barbed end^30, 31, 50^, curtailing its processivity. The F-bar domain containing protein Hof1, on the other hand, inhibits nucleation by formin^51^. In the future, it will be interesting to explore if other factors that bind either CP, formin, twinfilin or actin barbed ends, like CARMIL^16, 37^, V-1/myotrophin^37^, APC^39^, Spire^52^, IQGAP^53^ and polyphosphoinositides^40, 54^, might also form their own multiprotein ecosystems with these proteins at the barbed ends to tune actin assembly.

Where in a cell would the mechanisms uncovered here be relevant? Twinfilin, CP and formins operate simultaneously in a number of cellular compartments including filopodia, lamellipodia and stereocilia^6, 18, 21–24^ where size and elongation rates of actin filaments are tightly controlled. Given the high barbed-end affinity and cellular concentration of proteins like formins, CP, twinfilin and Ena/VASP, we expect that newly nucleated free barbed ends initiated by Arp2/3 branching would be rapidly captured by one of these factors. Filaments taken by formins would continue to grow and those captured by CP or twinfilin would stop growing and shrink at their barbed ends. Although formins are mainly thought of in the context of structures which consist of linear bundles, formins like FMNL2 have been shown to play a role in lamellipodial extension as well^24^. In light of our results, we speculate that the majority of growing actin filaments might be protected by formins or Ena/VASP bound ends. Although, it remains to be tested if twinfilin also favors Ena/VASP over CP and acts as a pro-Ena/VASP factor. Future cellular experiments where twinfilin activity can be locally enhanced/suppressed by optogenetic techniques are needed to test these predictions.

Living cells operate in a highly dynamic mechanical environment influenced by the extracellular matrix, neighboring cells and tissues. Formin is a mechanosensitive protein and its processivity is dramatically reduced under force^35^. In a number of cellular structures including filopodia and the cytokinetic ring, formin-based elongation is expected to occur under force e.g., the restoring force of the plasma membrane and contractile forces originating from actomyosin contraction. It will therefore be interesting for future studies to investigate the crosstalk between mechanical forces and biochemical factors, and how they co-regulate barbed-end assembly. It will also be important to examine if the novel interactions between twinfilin, CP and formin reported here might also play a role in human diseases and disorders including cancer invasion and progression^55, 56^, hearing loss^57^ as well as neuropathies and cardiac conditions^58^ in which these proteins have been implicated.

## Methods

### Purification and labeling of actin

Rabbit skeletal muscle actin was purified from acetone powder^59^ generated from frozen ground hind leg muscle tissue of young rabbits (PelFreez, USA). Lyophilized acetone powder stored at −80°C was mechanically sheared in a coffee grinder, resuspended in G-buffer (5 mM Tris-HCl pH 7.5, 0.5 mM Dithiothreitol (DTT), 0.2 mM ATP and 0.1 mM CaCl_2_), and cleared by centrifugation for 20 min at 50,000 × *g*. Supernatant was collected and further filtered with Whatman paper. Actin was then polymerized overnight at 4°C, slow stirring, by the addition of 2 mM MgCl_2_ and 50 mM NaCl to the filtrate. The next morning, NaCl powder was added to a final concentration of 0.6 M and stirring was continued for another 30 min at 4°C. Then, F-actin was pelleted by centrifugation for 150 min at 120,000 × *g*, the pellet was solubilized by dounce homogenization and dialyzed against G-buffer for 48 h at 4°C. Monomeric actin was then precleared at 435,000 × *g* and loaded onto a Sephacryl S-200 16/60 gel-filtration column (Cytiva, USA) equilibrated in G-Buffer. Fractions containing actin were stored at 4°C.

To biotinylate actin, purified G-actin was first dialyzed overnight at 4°C against G-buffer lacking DTT. The monomeric actin was then polymerized by addition of an equal volume of 2X labeling buffer (50 mM imidazole pH 7.5, 200 mM KCl, 0.3 mM ATP, 4 mM MgCl_2_). After 5 min, the actin was mixed with a 5-fold molar excess of NHS-XX-Biotin (Merck KGaA, Germany) and incubated in the dark for 15 h at 4°C. The F-actin was pelleted as above, and the pellet was rinsed with G-buffer, then homogenized with a dounce and dialyzed against G-buffer for 48 h at 4°C. Biotinylated monomeric actin was purified further on an Sephacryl S-200 16/60 gel-filtration column as above. Aliquots of biotin-actin were snap frozen in liquid N_2_ and stored at −80 °C.

To fluorescently label actin, G-actin was polymerized by dialyzing overnight against modified F-buffer (20 mM PIPES pH 6.9, 0.2 mM CaCl_2,_ 0.2 mM ATP, 100 mM KCl)^33^. F-actin was incubated for 2 h at room temperature with a 5-fold molar excess of Alexa-488 NHS ester dye (Thermo Fisher Scientific, USA). F-actin was then pelleted by centrifugation at 450,000 × *g* for 40 min at room temperature, and the pellet was resuspended in G-buffer, homogenized with a dounce and incubated on ice for 2 h to depolymerize the filaments. The monomeric actin was then re-polymerized on ice for 1 h by addition of 100 mM KCl and 1 mM MgCl_2_. F-actin was once again pelleted by centrifugation for 40 min at 450,000 × *g* at 4°C. The pellet was homogenized with a dounce and dialyzed overnight at 4°C against 1 L of G-buffer. The solution was precleared by centrifugation at 450,000 × *g* for 40 min at 4°C. The supernatant was collected, and the concentration and labeling efficiency of actin was determined.

### Purification and labeling of mTwf1 polypeptides

Wild type and mutant mouse mTwf1 plasmids were a gift from Pekka Lappalainen^20^. All of these proteins were expressed in *E. coli* BL21 (pRare). Cells were grown in Terrific Broth to log phase at 37°C. Expression was induced overnight at 18°C by addition of 1 mM IPTG. Cells were harvested by centrifugation at 11,200 × *g* for 15 min and the cell pellets were stored at −80°C. For purification, frozen pellets were thawed and resuspended in 35 mL lysis buffer (50 mM sodium phosphate buffer pH 8, 20 mM imidazole, 300 mM NaCl, 1 mM DTT, 1 mM PMSF and protease inhibitors (pepstatin A, antipain, leupeptin, aprotinin, and chymostatin, 0.5 μM each). Cells were lysed using a tip sonicator while being kept on ice. The cell lysate was then centrifuged at 120,000 × *g* for 45 min at 4°C. The supernatant was then incubated with 1 mL of Ni-NTA beads (Qiagen, USA) while rotating for 2 h at 4°C. The beads were then washed three times with the wash buffer (50 mM sodium phosphate buffer pH 8, 300 mM NaCl, 20 mM imidazole and 1 mM DTT). The beads were then transferred to a disposable column (Bio-Rad, USA). Protein was eluted using the elution buffer (50 mM phosphate buffer pH 8, 300 mM NaCl, 250 mM imidazole and 1 mM DTT). Fractions containing the protein were concentrated and loaded onto a size exclusion Superdex 75 Increase 10/300 column (Cytiva, USA) pre-equilibrated with 20 mM HEPES pH 7.5, 1 mM EDTA, 50 mM KCl and 1 mM DTT. Peak fractions were collected, concentrated, aliquoted, and flash-frozen in liquid N_2_ and stored at −80°C.

Mouse His-SNAP-mTwf1 plasmid was ordered from Twist biosciences. SNAP-mTwf1 was purified using the same protocol as above. Purified SNAP-mTwf1 was incubated with 5x excess of SNAP-surface-549 dye (New England Biolabs, Ipswich, MA) overnight at 4°C. Free dye was removed using a PD-10 desalting column (Cytiva, USA). Labeled protein was collected, concentrated, aliquoted, and flash-frozen in liquid N_2_ and stored at −80°C.

### Purification, labeling and biotinylation of formin mDia1

Mouse his-tagged mDia1 (FH1-FH2-C) formin was expressed in *E. coli*; BL21(DE3) pLysS cells. Cells were grown in Terrific Broth to log phase at 37°C. Expression was induced overnight at 18°C by addition of 1 mM IPTG. Cells were harvested by centrifugation at 11,200 × *g* for 15 min and the cell pellets were stored at −80°C. For purification, frozen pellets were thawed and resuspended in 35 mL lysis buffer (50 mM sodium phosphate buffer pH 8, 20 mM imidazole, 300 mM NaCl, 1 mM DTT, 1 mM PMSF and protease inhibitors (0.5 μM each of pepstatin A, antipain, leupeptin, aprotinin, and chymostatin). Cells were lysed using a tip sonicator while being kept on ice. The cell lysate was then centrifuged at 120,000 × *g* for 45 min at 4°C. The supernatant was then incubated with 1 mL of Ni-NTA beads (Qiagen, USA) while rotating for 2 h at 4°C. The beads were then washed three times with the wash buffer (50 mM sodium phosphate buffer pH 8, 300 mM NaCl, 20 mM imidazole and 1 mM DTT) and were then transferred to a disposable column (Bio-Rad, USA). Protein was eluted using the elution buffer (50 mM phosphate buffer pH 8, 300 mM NaCl, 250 mM imidazole and 1 mM DTT). Fractions containing the protein were concentrated and loaded onto a size exclusion Superdex 200 increase 10/300 GL column (Cytiva, USA) pre-equilibrated with 20 mM HEPES pH 7.5, 150 mM KCl, 10% glycerol and 0.5 mM DTT. Peak fractions were collected, concentrated, aliquoted, and flash-frozen in liquid N_2_ and stored at −80°C.

SNAP-mDia1^31^ was expressed and purified using the protocol described above. Purified SNAP-mDia1 was incubated with 5x excess of SNAP-surface-649 dye (New England Biolabs, USA) overnight at 4°C. Free dye was removed using a Superdex 200 increase 10/300 GL column (Cytiva, USA). Labeled protein was collected, concentrated, aliquoted, and flash-frozen in liquid N_2_ and stored at −80°C.

Biotin-SNAP-mDia1 prepared by incubating purified SNAP-mDia1 with Benzylguanine-Biotin (New England Biolabs, USA) according to the manufacturer’s instructions. Free biotin was removed using size-exclusion chromatography by loading the labeled protein on a Superose 6 gel-filtration column (GE Healthcare, Pittsburgh, PA) eluted with 20 mM HEPES pH 7.5, 150 mM KCl and 0.5 mM DTT.

### Purification and labeling of capping protein

Mouse his-tagged capping protein was expressed in *E. coli*; BL21(DE3) pLysS cells. Capping protein α1 and β2 were expressed from the same plasmid with a single His-tag on the alpha subunit^30^. Cells were grown in Terrific Broth to log phase at 37°C. Expression was induced overnight at 18°C by addition of 1 mM IPTG. Cells were harvested by centrifugation at 11, 200 × *g* for 15 min and the cell pellets were stored at −80°C. For purification, frozen pellets were thawed and resuspended in 35 mL lysis buffer (50 mM sodium phosphate buffer pH 8, 20 mM imidazole, 300 mM NaCl, 1 mM DTT, 1 mM PMSF and protease inhibitors (0.5 μM each of pepstatin A, antipain, leupeptin, aprotinin, and chymostatin). Cells were lysed using a tip sonicator while being kept on ice. The cell lysate was then centrifuged at 120,000 × *g* for 45 min at 4°C. The supernatant was incubated with 1 mL of Ni-NTA beads (Qiagen, USA) while rotating for 2 h at 4°C. The beads were then washed three times with the wash buffer (50 mM sodium phosphate buffer pH 8, 300 mM NaCl, 20 mM imidazole and 1 mM DTT) and transferred to a disposable column (Bio-Rad, USA). Protein was eluted using the elution buffer (50 mM phosphate buffer pH 8, 300 mM NaCl, 250 mM Imidazole and 1 mM DTT). Fractions containing the protein were concentrated and loaded onto a size exclusion Superdex75 Increase 10/300 column (Cytiva, USA) pre-equilibrated with 20 mM Tris-HCl, 50 mM KCl and 1 mM DTT. Peak fractions were collected, concentrated, aliquoted, and flash-frozen in liquid N_2_ and stored at −80°C.

SNAP-CP was expressed from a single plasmid containing His- and SNAP-tagged β1 subunit and untagged α1 subunit^31^. It was purified using the protocol described above. Purified SNAP-CP was incubated with 5x excess of SNAP-surface-549 dye or SNAP-surface-649 dye (New England Biolabs, USA) overnight at 4°C. Free dyes were removed using PD-10 desalting columns (Cytiva, USA). Labeled protein was collected, concentrated, aliquoted, and flash-frozen in liquid N_2_ and stored at −80°C.

### Purification of profilin

Human profilin-1 was expressed in *E. coli* strain BL21 (pRare) to log phase in LB broth at 37°C and induced with 1 mM IPTG for 3 h at 37°C. Cells were then harvested by centrifugation at 15,000 × *g* at 4°C and stored at −80°C. For purification, pellets were thawed and resuspended in 30 mL lysis buffer (50 mM Tris-HCl pH 8, 1 mM DTT, 1 mM PMSF protease inhibitors (0.5 μM each of pepstatin A, antipain, leupeptin, aprotinin, and chymostatin) was added, and the solution was sonicated on ice by a tip sonicator. The lysate was centrifuged for 45 min at 120,000 × *g* at 4°C. The supernatant was then passed over 20 ml of Poly-L-proline conjugated beads in a disposable column (Bio-Rad, USA). The beads were first washed at room temperature in wash buffer (10 mM Tris pH 8, 150 mM NaCl, 1 mM EDTA and 1 mM DTT) and then washed again with 2 column volumes of 10 mM Tris pH 8, 150 mM NaCl, 1 mM EDTA, 1 mM DTT and 3 M urea. Protein was then eluted with 5 column volumes of 10 mM Tris pH 8, 150 mM NaCl, 1 mM EDTA, 1 mM DTT and 8 M urea. Pooled and concentrated fractions were then dialyzed in 4 L of 2 mM Tris pH 8, 0.2 mM EGTA, 1 mM DTT, and 0.01% NaN_3_ (dialysis buffer) for 4 h at 4°C. The dialysis buffer was replaced with fresh 4 L buffer and the dialysis was continued overnight at 4°C. The protein was centrifuged for 45 min at 450,000 × *g* at 4°C, concentrated, aliquoted, flash frozen in liquid N_2_ and stored at −80°C.

### Conventional TIRF microscopy for single molecule imaging

Glass coverslips (60 x 24 mm; Thermo Fisher Scientific, USA) were first cleaned by sonication in detergent for 60 min, followed by successive sonications in 1 M KOH and 1 M HCl for 20 min each, and in ethanol for 60 min. Coverslips were then washed extensively with H_2_O and dried in an N_2_ stream. The cleaned coverslips were coated with 2 mg/mL methoxy-polyethylene glycol (mPEG)-silane MW 2,000 and 2 µg/mL biotin-PEG-silane MW 3,400 (Laysan Bio, USA) in 80% ethanol (pH 2.0) and incubated overnight at 70°C. Flow cells were assembled by rinsing PEG-coated coverslips with water, drying with N_2_, and adhering to μ-Slide VI0.1 (0.1 x 17 x 1 mm) flow chambers (Ibidi, Germany) with double-sided tape (2.5 cm x 2 mm x 120 μm) and 5 min epoxy resin (Devcon, USA). Before each reaction, the flow cell was sequentially incubated for 1 min each with 4 μg/ml streptavidin and 1% BSA in 20 mM HEPES pH 7.5, and 50 mM KCl. The flow cell was then equilibrated with TIRF buffer (10 mM imidazole, pH 7.4, 50 mM KCl, 1 mM MgCl_2_, 1 mM EGTA, 0.2 mM ATP, 10 mM DTT, 2 mM DABCO and 0.5% methylcellulose [4,000 cP]). 1 µM G-actin and 0.5 µM Profilin along with 100 pM 649-mDia1 in TIRF buffer were introduced into the flow cell and filaments were allowed to grow for 2 to 3 min. The flow cell was then rinsed with TIRF buffer to remove free formins, and the solution was replaced with 1 µM G-actin and 4 µM profilin and 10 nM 549-CP (with or without mTwf1). In experiments with fluorescently labeled twinfilin, 10 nM unlabeled CP along with 40 nM 549-mTw1 was introduced into the flow cell. Each experiment was repeated at least three times. Data from a single replicate is presented in the figures.

### Microfluidics-assisted TIRF (mf-TIRF) imaging and analysis

Actin filaments were first assembled in microfluidics-assisted TIRF (mf-TIRF) flow cells^33, 34^. Coverslips were first cleaned by sonication in Micro90 detergent for 20 min, followed by successive 20 min sonications in 1 M KOH and 1 M HCl for 20 min each and 200 proof ethanol. Washed coverslips were then stored in fresh 200 proof ethanol. Coverslips were then washed extensively with H_2_O and dried in an N_2_ stream. These dried coverslips were coated with 2 mg/mL methoxy-poly(ethylene glycol) (mPEG)-silane MW 2,000 and 2 µg/mL biotin-PEG-silane MW 3,400 (Laysan Bio, USA) in 80% ethanol (pH 2.0) and incubated overnight at 70°C. A 40 µm high PDMS mold with 3 inlets and 1 outlet was mechanically clamped onto a PEG-Silane coated coverslip. The chamber was then connected to a Maesflo microfluidic flow-control system (Fluigent, France), rinsed with mf-TIRF buffer (10 mM imidazole, pH 7.4, 50 mM KCl, 1 mM MgCl_2_, 1 mM EGTA, 0.2 mM ATP, 10 mM DTT, 1 mM DABCO) and incubated with 1% BSA and 10 µg/mL streptavidin in in 20 mM HEPES pH 7.5, and 50 mM KCl for 5 min. 100 pM Biotin-SNAP-mDia1 molecules in TIRF buffer were then flowed in and allowed to anchor on the glass coverslip. Actin filaments with free pointed ends and barbed ends anchored to the formins were grown by flowing in a solution containing 1µM G-actin (15% Alexa-488 labeled) and 0.5 µM profilin in mf-TIRF buffer. All experiments were carried out at room temperature in TIRF buffer. Each experiment was repeated at least three times. Data from a single replicate is presented in the figures.

Single-wavelength time-lapse TIRF imaging was performed on a Nikon-Ti2000 inverted microscope equipped with a 40 mW Argon laser, a 60X TIRF-objective with a numerical aperture of 1.49 (Nikon Instruments Inc., USA) and an IXON LIFE 888 EMCCD camera (Andor Ixon, UK). One pixel was equivalent to 144 × 144 nm. Focus was maintained by the Perfect Focus system (Nikon Instruments Inc., Japan). Time-lapsed images were acquired every 2 s or 5 s using Nikon Elements imaging software (Nikon Instruments Inc., Japan).

Images were analyzed in Fiji^60^. Background subtraction was conducted using the rolling ball background subtraction algorithm (ball radius 5 pixels). Time-lapse images of between 50 and 100 filaments were acquired in a single field of view for each condition and all of these filaments were included to determine the cumulative distribution functions (CDFs) showing the time-dependent survival fraction of various complexes. For mf-TIRF assays, the kymograph plugin was used to draw kymographs of individual filaments. The kymographs were used to identify the timepoint of detachment of filaments as a function of time. Data analysis and curve fitting were carried out in Microcal Origin.

### Single-molecule imaging and analysis

Glass coverslips (60 x 24 mm; Thermo Fisher Scientific, USA) first cleaned by sonication in Micro90 detergent for 20 min, followed by successive 20 min sonications in 1 M KOH and 1 M HCl for 20 min each and 200 proof ethanol. Washed coverslips were then stored in fresh 200 proof ethanol. Coverslips were then washed extensively with H_2_O and dried in an N_2_ stream. The cleaned coverslips were coated with 2 mg/mL methoxy-poly(ethylene glycol) (mPEG)-silane MW 2,000 and 2 µg/mL biotin-PEG-silane MW 3,400 (Laysan Bio, USA) in 80% ethanol (pH 2.0) and incubated overnight at 70°C. Flow cells were assembled by rinsing PEG-coated coverslips with water, drying with N_2_, and adhering to μ-Slide VI0.1 (0.1 x 17 x 1 mm) flow chambers (Ibidi, Germany) with double-sided tape (2.5 cm x 2 mm x 120 μm) and 5-min epoxy resin (Devcon, USA). Before each reaction, the flow cell was sequentially incubated for 1 min each with 4 μg/ml streptavidin in TIRF buffer (10 mM imidazole pH 7.4, 50 mM KCl, 1 mM MgCl_2_, 1 mM EGTA, 0.2 mM ATP, 10 mM DTT, 1 mM DABCO and 0.5% methylcellulose [4,000 cP]) and 1% BSA in TIRF buffer. Actin filaments were grown using 0.5 µM G-actin (15% Alexa-488 labeled and 0.5% biotinylated G-actin), 1 µM profilin and 50 pM 649-mDia1. The filaments were allowed to elongate for a few minutes. The chamber was then rinsed with TIRF buffer and a solution containing 10 nM CP (unlabeled or 549-CP) and/or mTwf1 (unlabeled or 549-mTwf1) in presence of 0.5 µM G-actin (15% Alexa-488 labeled and 0.5% biotinylated G-actin) and 1 µM profilin was introduced into the flow cell. The flow cell was then placed on a Nikon-Ti2-E inverted microscope equipped with a 60X TIRF-objective with a numerical aperture of 1.49 (Nikon Instruments Inc., USA) and an IXON LIFE 888 EMCCD camera (Andor Ixon, UK). The sample was sequentially excited by 488 nm, 561 nm and 640 nm lasers and imaged on the EMCCD camera. Images were acquired either continuously or with 4 s delay between consecutive images. All experiments were carried out at room temperature in TIRF buffer. Each experiment was repeated at least three times.

For three color experiments involving 549-mTwf1, 649-CapZ, and 488-actin, actin filaments were elongated from 1 µM G-actin (15% Alexa-488 labeled and 1.4% biotinylated G-actin). Free actin monomers were removed by rinsing the flow cell with excess of TIRF buffer and the filaments were exposed to a solution containing 10 nM 649-CP. Following capping, the chamber was once again rinsed with TIRF buffer to remove free 649-CP and then 15 nM 549-mTwf1 was introduced. The sample was sequentially excited by 488 nm, 561 nm and 640 nm lasers. The images were captured on the EMCCD camera and acquired with 1 s delay between consecutive images.

The three-color images thus acquired were then analyzed in Fiji^60^. A 5 × 5-pixel box was drawn at the location of the barbed end of the filament and the time-dependent integrated intensity values were recorded for the single-molecule channels. The integrated intensity values were background corrected by subtracting integrated intensity of a 5 x 5 pixel box drawn away from the filament.

### Determination of rates of CP or formin dissociation from BFC complexes (*k’_-F_* and *k’_-C_)*

The time-dependent fraction of BFC complexes dissociating by either transitioning to BF (re-elongation) or BC (detachment) upon addition of twinfilin to the flow were plotted versus time (black symbols in Fig. 2e). The kinetics of dissociation of BFC and simultaneous appearance of BF (magenta symbols) and BC (yellow symbols) were analyzed using the approach described in our earlier study ^30^. Briefly, BFC dissociation can occur via one of the following two routes:

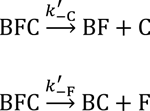

These reactions can be described by the following differential equations:

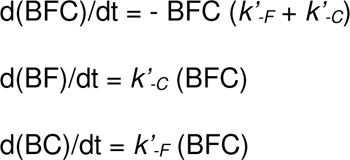

The number of filaments transitioning out of BFC, and into BF and BC all vary exponentially with rate constant (*k’_-F_* + *k’_-C_*). The value of *k’_-F_* and *k’_-C_* was derived from the relative fraction of filaments in BF and BC states as follows.

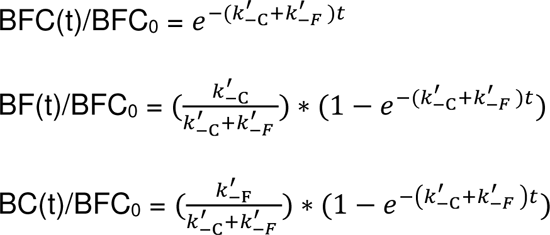

BFC_0_ is the total number of filaments in BFC state just prior to flowing in twinfilin. The ratio of number of filaments taking either of the two routes (BF or BC) is given by N_BF_/N_BC_ = *k’_-C_*/*k’_-F_*. Note that *k’_-F_* + *k’_-C_* are observed rates as they depend upon the concentration of twinfilin as well as elongation rate of formins.

### Statistical analysis and error bars for dissociation rates

The uncertainty in dissociation rates of formin (BF→B + F) or BFC decision complexes (BFC → BF +C or BFC → BC + F) were determined by bootstrapping strategy^35^. The dissociation rate was determined by fitting the survival fraction (or CDF) data to a single exponential function (y = e^-*k*t^ or y = 1 – e^-*k*t^). A custom-written MATLAB code was then used to simulate BF (or BFC) complex lifetimes for N filaments (where N is the number of filaments in the particular experiment) based on the rate k determined from the experimental data. The simulation was repeated 1000 times to generate 1000 individual survival fractions of N filaments. Each dataset was then fit to an exponential function and a rate *k_obs_* was determined for each of the 1000 simulated datasets. The standard deviation of these estimated rates allowed us to determine the uncertainty in our measured rates. Statistical comparison between indicated conditions was conducted using two-sample t-test. Differences were considered significant if the P value was <0.05.

## Data availability

Data supporting the findings of this manuscript are available from the corresponding author upon reasonable request.

## Acknowledgements

We thank Pekka Lappalainen for generously sharing plasmids for wild type and mutant mTwf1 as well as for advice on SNAP-tagging of mTwf1. We thank Jim Bear for fruitful discussions and feedback on the manuscript. We thank Ankita for helping with custom MATLAB codes for statistical analysis and for comments on the manuscript. We thank Jan Faix for the gift of VASP chimera constructs. Shashank Shekhar thanks Marie-France Carlier, Bruce Goode and Jeff Gelles for many years of inspiring discussions on barbed-end dynamics. This work was supported by NIH NIGMS grant R35GM143050 to Shashank Shekhar and startup funds from Emory University.

## Author contributions

HU and IG conducted experiments, HU and IG analyzed data, HU and SS prepared figures. SS designed experiments and supervised the project. SS and HU wrote the first draft of the manuscript and all authors contributed to the editing. SS acquired funding.

## Competing interests

We declare no conflicts of interest.

**Correspondence** and requests for materials should be addressed to S.S.

