## Supplementary Info for "Multicomponent regulation of actin barbed end assembly by twinfilin, formin and capping protein"

**This PDF file includes:**

Supplementary Figs. 1-6, and their captions.

Captions for Supplementary Movies 1-4

**Supplementary Fig. 1: Presence of 1 µM mTwf1 does not change the elongation rate of formin mDia1.**

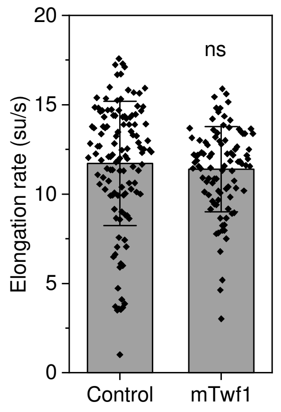


Rates (± sd) of barbed end elongation of formin-anchored filaments in presence of 0.2 µM unlabeled G-actin and 0.7 µM profilin in absence or presence of 1 µM mTwf1. Column height is indicates the mean, and whiskers indicate sd. Statistical differences: ns (no evidence for significance at $p$ = 0.05) using two-sample t test. Number of filament ends analyzed for each condition (left to right): 123, 96. All experiments were performed at least three independent times, and yielded similar results. Data shown are from one experiment.

**Supplementary Fig. 2: Concentration dependent effect of mTwf1 on dissociation rate of BFC complexes and appearance of BF and BC.**

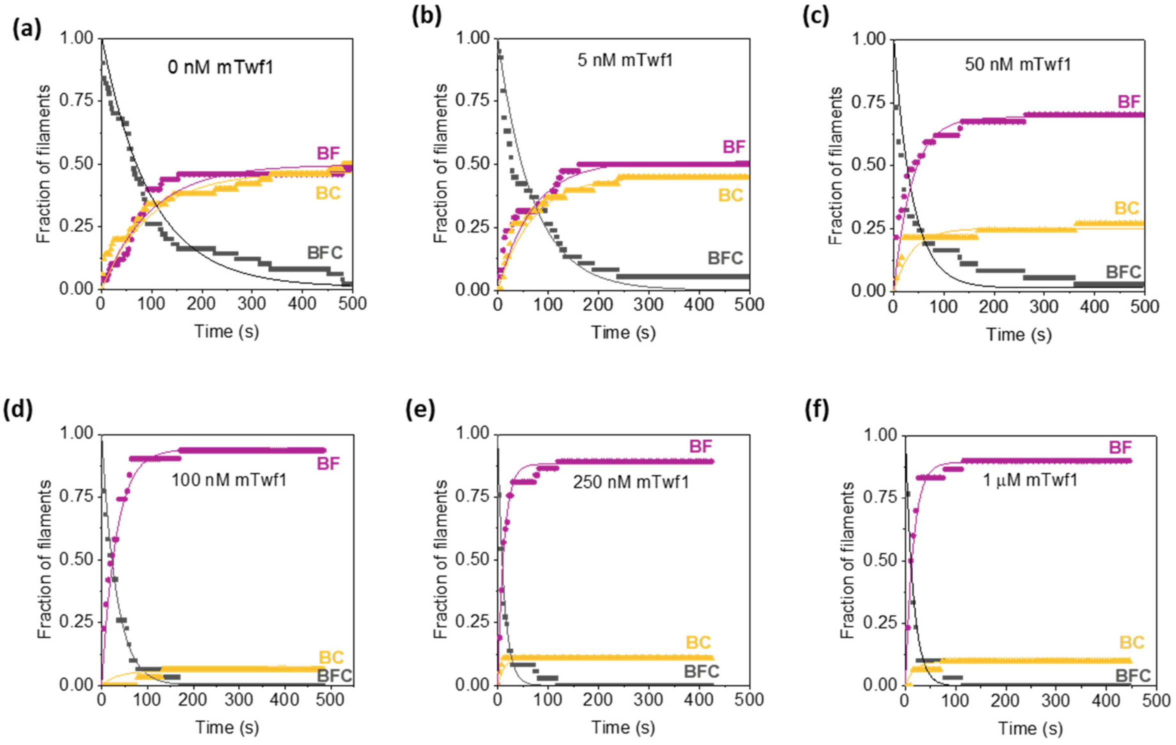


Actin filaments were nucleated from coverslip-anchored formins by introducing a flow containing 1 µM G-actin (15% Alexa-488 labeled) and 0.5 µM profilin. The filaments were then exposed to a flow containing 1 µM unlabeled G-actin, 4 µM profilin and 1 µM CP for about 10 s to convert formin-bound barbed ends (BF) to formin-CP bound barbed ends or "decision complexes" (BF + C →BFC). These BFC complexes were then exposed to a flow containing PA (control) only or supplemented with varying concentrations of mTwf1. **(a)** Control (n=50 filaments). Survival fraction of BFC (black symbols) filaments transitioning either to BF (filaments resuming elongation, magenta symbols) or BC (filaments detached, yellow symbols). The experimental data (symbols) is fitted to exponential fits (lines), such that $k_{-BFC} = k'_{-F} + k'_{-C}$, where $k'_{-F}$ is the dissociation rate of formin from BFC (BFC → BC + F) and $k'_{-C}$ is the dissociation rate of CP from BFC (BFC → BF + C) (see methods). **(b)** same as (a) but for 5 nM mTwf1 (n=38 filaments) **(c)** same as (a) but for 50 nM mTwf1 (n=37 filaments) **(d)** same as (a) but for 100 nM mTwf1 (n=31 filaments) **(e)** same as (a) but for 250 nM mTwf1 (n=37 filaments) **(f)** same as (a) but for 1 µM mTwf1 (n=30 filaments). All experiments were performed at least three independent times, and yielded similar results. Data shown are from one experiment.

**Supplementary Fig. 3: Photobleaching tests of 549-CP molecules.**

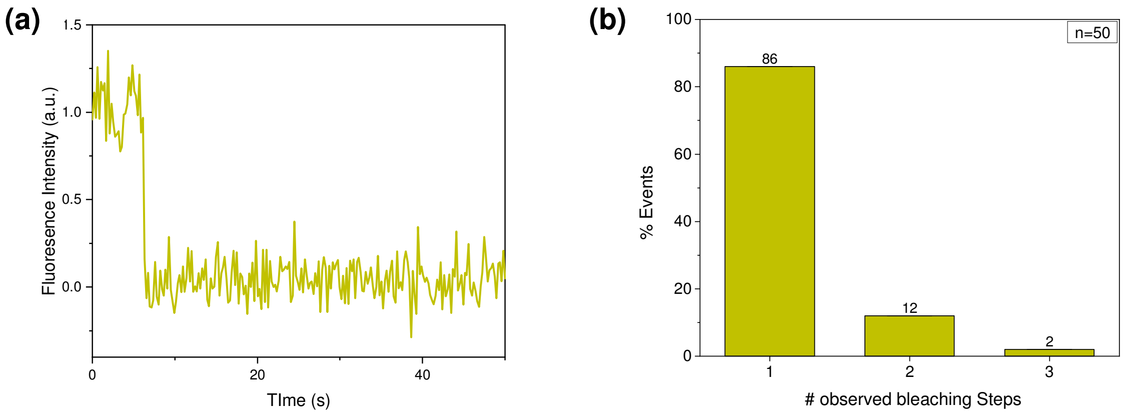


**(a)** An example of a single-step bleaching profile of a 549-CP molecule. **(b)** Histogram showing the number of steps leading to complete bleaching of individual fluorescent spots. A total of 50 spots were monitored. Photobleaching data confirmed that majority of 549-CP molecules were labelled with only one dye molecule, consistent with a single SNAP tag per heterodimeric CP molecule.

**Supplementary Fig. 4: Photobleaching tests of 649-mDia1 molecules.**

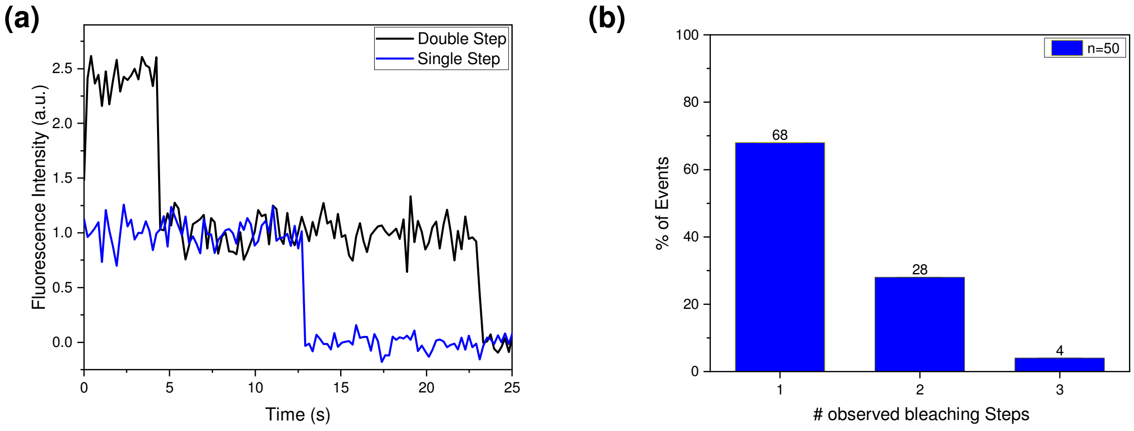


**(a)** An example of a single-step (blue curve) and double step (black curve) bleaching profile of two separate 649-mDia1 molecules. **(b)** Histogram showing the number of steps leading to complete bleaching of individual fluorescent spots. A total of 50 spots were monitored. Photobleaching tests of 649-mDia1 showed that majority of these molecules exhibited a single- or double-step bleaching profile, consistent with the dimeric nature of formin mDia1 molecules.

**Supplementary Fig. 5: SNAP tagging of mTwf1 doesn't influence its uncapping activities.**

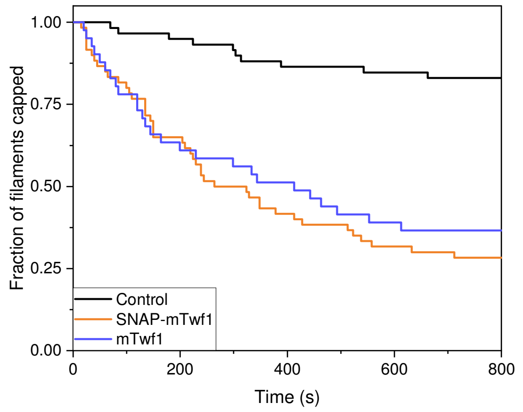


Pre-formed actin filaments (1 µM G-actin, 15% Alexa-488 labeled) were introduced into the mf-TIRF chamber and captured at their barbed ends by coverslip-anchored CP. These CP-bound filaments were then exposed to TIRF buffer only (control) or in presence of either 1 µM SNAP-mTwf1 (orange) or 1 µM mTwf1 (blue). Disappearance of actin filaments from the field of view as a function of time was recorded to quantify uncapping. No significant difference between uncapping activities of SNAP-mTwf1 and mTwf1 was observed.

**Supplementary Fig. 6: Photobleaching tests of 549-mTwf1 molecules**

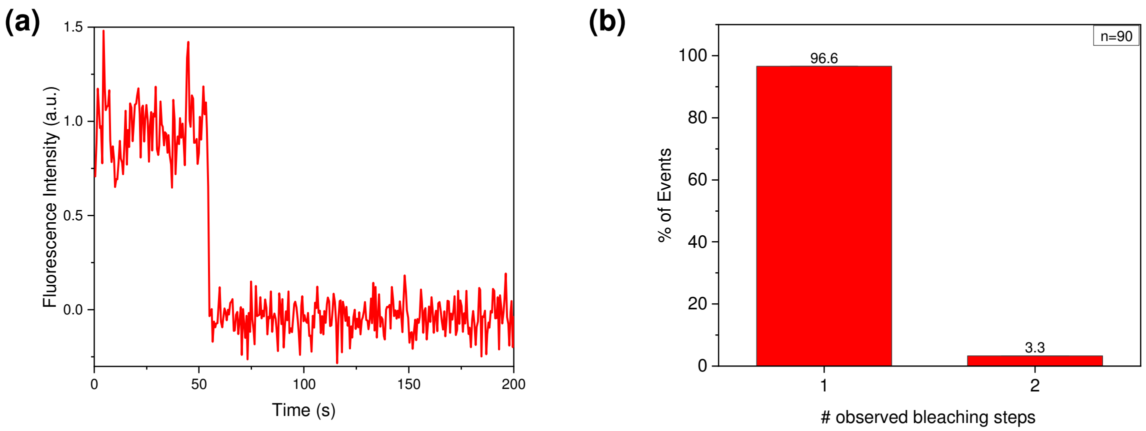


**(a)** An example of a single-step bleaching profile of a 549-mTwf1 molecule. **(b)** Histogram showing the number of steps leading to complete bleaching of individual fluorescent spots. A total of 90 spots were monitored. Photobleaching data confirmed that majority of 549-mTwf1 molecules were labelled with only one dye molecule, consistent with a single SNAP tag per mTwf1 molecule.

### Movie Legends

**Supplementary Movie 1. Elongation of formin-anchored filaments in presence of profilin and actin.**

Actin filaments were nucleated from coverslip-anchored formins by introducing a flow containing 1 µM G-actin (15% Alexa-488 labeled) and 0.5 µM profilin. The fluorescent filaments were then elongated in presence of 1 µM unlabeled G-actin and 4 µM profilin to ensure insertional elongation between fluorescent fragment and surface-anchored formins. These filaments were then exposed to a flow containing 0.2 µM unlabeled G-actin and 0.7 µM profilin (also see figure 1b).

**Supplementary Movie 2. Elongation of formin-anchored filaments in presence of CP, profilin and actin.**

Actin filaments were nucleated from coverslip-anchored formins by introducing a flow containing 1 µM G-actin (15% Alexa-488 labeled) and 0.5 µM profilin. The fluorescent filaments were then elongated in presence of 1 µM unlabeled G-actin and 4 µM profilin to ensure insertional elongation between fluorescent fragment and surface-anchored formins. These filaments were then exposed to a flow containing 0.2 µM unlabeled G-actin, 0.7 µM profilin and 50 nM CP (also see figure 1c).

**Supplementary Movie 3. Elongation of formin-anchored filaments in presence of CP, twinfilin, profilin and actin.**

Actin filaments were nucleated from coverslip-anchored formins by introducing a flow containing 1 µM G-actin (15% Alexa-488 labeled) and 0.5 µM profilin. The fluorescent filaments were then elongated in presence of 1 µM unlabeled G-actin and 4 µM profilin to ensure insertional elongation between fluorescent fragment and surface-anchored formins. These filaments were then exposed to a flow containing 0.2 µM unlabeled G-actin, 0.7 µM profilin, 50 nM CP and 1 µM mTwf1 (also see figure 1e).

**Supplementary Movie 4. Direct visualization of Formin-CP decision complex dynamics by single molecule imaging.**

Multicolor, merged, single molecule time-lapse movie of an actin filament (cyan) elongating with 649-mDia1 (magenta) bound to its barbed end in presence of 0.5 µM G-actin (15% Alexa-488 labeled, 0.5% biotin labeled) and 1 µM profilin and 10 nM 549-CP (yellow). The magenta arrowhead denotes the location of translocating formin, white arrowhead denotes the formation of the 549-CP:649-mDia1 decision complex and the yellow arrowhead marks the departure of 649-mDia1 from the decision complex, leaving 549-CP behind (also see figure 3a-c).
